# Investigating Cell Viability under Shear Stress in Complex Microstreaming Flows Generated by Ultrasound-Driven Actuated Microbubbles

**DOI:** 10.1101/2024.09.27.615272

**Authors:** Amirabas Bakhtiari, Benedikt Schumm, Martin Schönfelder, Christian J. Kähler

**Affiliations:** Institute for Fluid Mechanics and Aerodynamics, University of the Bundeswehr Munich Werner-Heisenberg-Weg 39, 85579 Neubiberg, Germany; Professorship of Exercise Biology, Department of Health and Sport Sciences, Technische Universität München, 80809 Munich, Germany

**Keywords:** Cell Viability, Cell Manipulation, Shear Stress, Acoustofluidics, Microbubble Streaming

## Abstract

The analysis of rare or specialized cells is often a time-consuming process due to their low concentrations. In this study, we applied, for the first time, a method previously used on polymer particles to manipulate human cells. This technique enables the automatic direction and collection of target cells passing through a microchannel, significantly increasing their concentration for further analysis. The movement of the cells is controlled by an acoustically induced vortex flow generated by a microbubble. By modulating the activation of this microstreaming, the cells are shifted either to the upper or lower regions of the channel and directed into a side channel for collection downstream. The localized stress distribution, along with long-term testing that showed no cell damage, confirmed the biocompatibility of this method, making it a promising tool for lab-on-a-chip systems and biomedical diagnostics.

**Impact Statement:** This study presents an innovative use of ultrasound-driven microbubble streaming for the precise manipulation and sorting of human cells in microfluidic environments, all while maintaining cell viability. The research shows that the localized shear stress near the microbubble is significantly below the damage threshold for cells, confirming the biocompatibility of this method. The potential impact of this work is considerable for lab-on-a-chip systems and biomedical diagnostics. It offers a reliable, non-invasive solution for the manipulation, sorting, and removal of compromised cells, thus streamlining research and diagnostic procedures. By ensuring the safe and efficient handling of rare or specialized cells, this technique can accelerate various biomedical applications. Additionally, the study’s evidence of sustained cell viability under microstreaming conditions suggests broader applicability in biomedical devices, particularly in automated dead cell removal and selective cell positioning.

## 1. Introduction

Microfluidics is a rapidly advancing field that plays a crucial role in various biomedical and biological applications, particularly those requiring precise manipulation of particles or cells within micro-scale environments. This technology leverages different force fields, such as acoustic, electric, magnetic, and optical forces, to achieve contactless manipulation of microscopic entities. The ability to control cells and particles with high precision has enabled significant advancements in lab-on-a-chip devices, which have been used for applications such as detection, sorting, mixing, and analysis at the single-cell level (Nilsson & Evander, 2009; Nan et al., 2014; Sheng et al., 2014). These applications are vital in areas such as circulating tumor cell (CTC) enrichment, hematopoietic stem cell (HSC) isolation, and single-cell electroporation, highlighting the importance of microfluidics in modern biomedical research (Armstrong et al., 2011; Bischoff et al., 2003; Khine et al., 2005).

Nonetheless, manipulating cells within these systems presents considerable challenges. It is essential to handle cells in a way that preserves their viability and functionality. For instance, using high-power lasers in optical tweezers or strong electric fields in electrokinetic tweezers can damage cell membranes or disrupt the experimental environment (Shields et al., 2015). Additionally, exposing cells to harsh conditions, such as excessive shear stress, can result in cell membrane damage, leading to cell death or altered behavior. These challenges highlight the need for biocompatible manipulation techniques that minimize the risk of cell damage during experimentation.

In the realm of microfluidics, various force fields have been explored for manipulating fluids and cells at the microscale. Among these, microbubble streaming—produced by acoustically actuated microbubbles—has emerged as a particularly promising technique due to its non-invasive and biocompatible characteristics. This method generates localized fluid flows that can be precisely controlled by modulating the driving frequency and amplitude, making it highly adaptable to different microchannel configurations and suitable for a broad spectrum of applications (Riley, 2001; Marmottant & Hilgenfeldt, 2003; Versluis et al., 2010). Over the past ten years, microbubble streaming has been effectively integrated into sophisticated lab-on-a-chip devices, facilitating critical functions such as particle sorting, fluid mixing, particle focusing, and size-selective separation. Additionally, it has proven valuable in mitigating clogging issues within microfluidic systems, underscoring its versatility and effectiveness (Thameem et al., 2016; Wang et al., 2013; Wang et al., 2010; Bakhtiari & Kähler, 2022; Bakhtiari & Kähler, 2023; Bakhtiari & Kähler, 2024; Bakhtiari & Kähler, 2024).

Despite its growing use, comprehensive studies that thoroughly evaluate the biocompatibility of microbubble streaming—particularly regarding its effects on cell viability under the shear stress it induces—are still lacking. Previous research has suggested that the outcomes of exposure to shear stress can vary widely, ranging from beneficial to potentially harmful, depending on factors such as the intensity and duration of the applied stress. For example, controlled shear stress has been shown to promote cellular proliferation and differentiation, which is beneficial in tissue engineering contexts (Baeyens et al., 2016). On the other hand, excessive shear stress can result in cellular damage or even apoptosis, especially in tissues that are delicate or compromised by disease (Koutsiaris et al., 2007; Roux et al., 2020). Research shows that in humans, normal physiological blood flow generates shear stress ranging from 0.1 to 9.5 Pa (Baeyens et al., 2016; Koutsiaris et al., 2007; Roux et al., 2020; Espina et al., 2023). This provides a reference for the natural shear stress environment cells typically experience, offering insight into what conditions are safe for maintaining cell integrity during experimentation.

In this study, we seek to address this gap in knowledge by experimentally investigating the effects of microbubble streaming on blood cells. Our research begins with the precise measurement of shear stress generated by microbubble actuation, with particular attention to the regions closest to the bubble surface where shear stress levels are highest. We employ high-frequency micro-particle tracking velocimetry (micro-PTV) to capture the shear stress distribution with high spatial resolution. Following this, we subject blood cells to these microbubble-induced flows and monitor their viability and behavior over time, paying close attention to any changes that may indicate cellular stress or damage. Additionally, we leverage automated cell positioning techniques driven by microbubble streaming to detect, track, and selectively remove dead cells from the fluid, ensuring that only healthy, viable cells remain within the system. This comprehensive study aims to provide vital insights into the biocompatibility of microbubble streaming, ultimately contributing to its safe and effective implementation in various microfluidic applications.

## 2. Experimental setup

In this section, we outline the experimental design illustrated in Figure 1. The experimental setup primarily comprises a microfluidic system, an optical configuration, and a control system.

**Figure 1.**
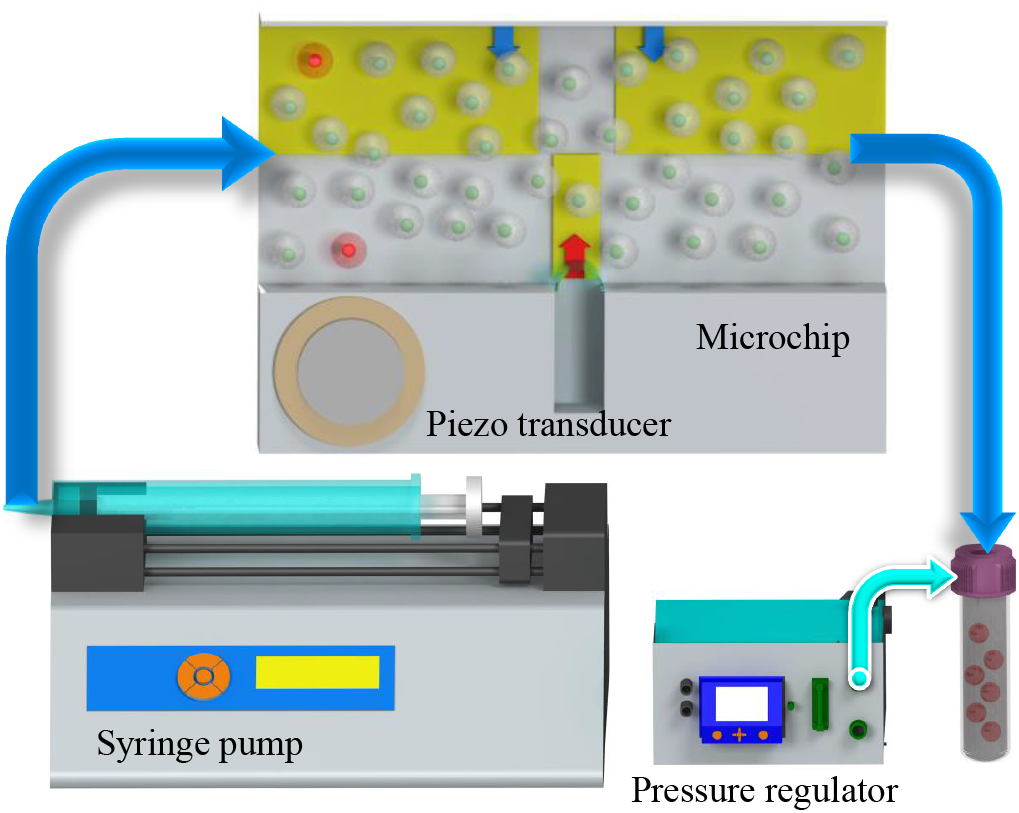
The microfluidic chip and its associated flow control system. The 20 mm microchannel features a rectangular cross-section (H = 100 μm × W = 500 μm), with a central cavity measuring w = 80 μm in width and h = 500 μm in length. This specific design allows for the controlled introduction and confinement of a gas pocket within the cavity when liquid is infused into the channel. The flow rate is regulated by a syringe pump, and a piezoelectric transducer stimulates the microbubble, while a pressure regulator ensures the stabilization of both liquid and bubble pressures.

### 2.1. Preparation of Samples

#### 2.1.1. Preparation of Polystyrene Microsphere Suspension for Shear Stress Measurement

To characterize the flow field of microbubble streaming and to determine the shear stress, polystyrene particles with a diameter of 2 μm (PS-FluoRed: Excitation/Emission 530 nm/607 nm) were employed as tracer particles. These particles were selected due to their low Stokes number, ensuring they accurately follow the streamlines. The polystyrene microspheres, stabilized with negatively charged sulfate groups to prevent agglomeration and adhesion, were procured from Microparticles GmbH, Germany. During the sample preparation process, the microspheres were suspended in an aqueous solution containing 23.8 w – w% glycerol in distilled water (Volk & Kähler, 2018; Volk & Kähler, 2018). This solution was prepared to achieve neutral buoyancy of the particles, effectively matching their density with that of the surrounding medium.

#### 2.1.2. Peripheral Blood Mononuclear Cells (PBMC) and Propidium Iodide Staining

Peripheral Blood Mononuclear Cells (PBMCs) were isolated from whole blood samples obtained from a male healthy donor (one of the authors) in accordance with institutional ethical guidelines. Blood samples were collected in EDTA monovettes (Sarstedt) to prevent coagulation and processed for cell isolation. The samples were diluted 1:2 with phosphate-buffered saline (PBS, pH 7.4, Gibco). Diluted blood (10 mL) was subjected to density gradient centrifugation using 5 mL Ficoll-Paque PLUS (density 1.077 g/mL; GE Healthcare). Blood samples were centrifuged at 400 × g for 30 minutes, at room temperature, with low acceleration, and no braking. After centrifugation, plasma was carefully removed, and buffy coat - containing PBMCs - was carefully collected, washed twice with PBS, and resuspended in PBS at a concentration of 1000 cells/μl.

To assess cell viability, propidium iodide (PI, Sigma-Aldrich) was used as a fluorescent stain for compromised cells. PI was added to the cell suspension at a final concentration of 2 μg/ml (∼ 3 μM) and incubated for 10 minutes at room temperature, protected from light. PI selectively binds to the DNA/RNA of dead cells with damaged membranes, emitting red fluorescence under excitation at 535 nm. The cell suspension, at a concentration of 1000 cells/μl, was prepared for microfluidic experiments to enable the automatic removal of dead cells and the monitoring of cell viability under shear stress induced by microbubble streaming. Approximately 0.1% of cells were damaged and detectable via fluorescence microscopy

### 2.2. Microfluidic System Configuration

The microfluidic system, as depicted in Figure 1, integrates transparent microchannels with sophisticated flow control mechanisms. The fabrication of the microchannels was carried out using the well-established soft lithography process, following the methodology outlined by Wang et al. (2012). The fabricated microchannel spans 20 mm in length and possesses a rectangular cross-section, with a height (*H*) of 100 μm and a width (*W*) of 500 μm. Centrally located within the channel is a cavity, measuring 80 μm in width (*w*) and 500 μm in length (*h*). This specific design allows for the controlled introduction and confinement of a gas pocket within the cavity when a liquid is infused into the channel.

The introduction of liquid into the microchannel results in the entrapment of a gas pocket, typically air, within the lateral cavity. This process leads to the formation of a quasi-cylindrical microbubble. It is noteworthy that the gas composition within the bubble is not limited to air; alternatives such as nitrogen, argon, and helium, can be utilized. The choice of gas can be crucial, particularly in scenarios where the interaction between the fluid and atmospheric air must be avoided to maintain the integrity of the conveyed fluids or particles.

When the microbubble is subjected to bulk acoustic waves, generated by a piezoelectric transducer operating at or near the bubble’s resonant frequency, it induces primary oscillatory flow within the channel. This primary flow subsequently gives rise to secondary flow patterns, characterized by counter-rotating vortices along the channel walls (Riley, 2001; Marmottant & Hilgenfeldt, 2003; Versluis et al., 2010).

To ensure experimental reproducibility, it is essential to maintain a consistent microbubble size. This can be achieved by continuously adjusting and controlling the pressure difference between the channel’s interior and the surrounding ambient pressure, as recommended by Volk et al. (2015). In this study, this approach was also employed to maintain a constant microbubble size throughout the experiments. This was achieved by precisely regulating the flow of the aqueous sample solution into the channel, using a syringe pump (neMESYS, Germany) and a pressure regulator (Fluigent MFCS™-EZ, 0–1000 mbar, France). Figure 1 provides a schematic overview of the experimental setup. For further elaboration on the setup and its components, readers are referred to Bakhtiari & Kähler (2022).

### 2.3. Optical setup

The optical setup consists of an upright Zeiss AxioImager.Z2 microscope equipped with a dichroic filter, 20 × objective lens (EC Plan Neouar 20× /0.50 M27), an sCMOS camera (pco.edge 5.5), and a Phantom v2640 ONYX high-speed camera, enabling both brightfield and darkfield imaging modalities.

Although both imaging modes are accessible in our system, darkfield imaging is particularly advantageous for monitoring cellular conditions. In this configuration, cells stained with propidium iodide (PI) become fluorescently detectable when their membranes are compromised. PI, a fluorescent dye, intercalates into the DNA of damaged cells and emits a red fluorescence upon excitation by a 520 nm wavelength high-power light-emitting diode (Prizmatix UHP-T-520-EP LED). The dichroic filter in the Zeiss AxioImager.Z2 selectively transmits the emitted fluorescence to the sCMOS camera while blocking the excitation light, enhancing the signal-to-noise ratio. This makes darkfield imaging highly effective for detecting compromised cells and filtering out undamaged cells, as well as eliminating unwanted artifacts such as fluid impurities or channel contamination during imaging. To avoid photo bleaching due to constant illumination, the light source was used in pulsed mode and synchronized with the camera.

Conversely, brightfield imaging, utilizing a 100 W halogen lamp, was employed for higher frame rate capture with the high-speed camera. The frame rate used was 3000 kHz. This approach is more effective for resolving shear stress in regions of rapid streaming near the bubble’s surface. Darkfield imaging is less suitable in this context, as the high-intensity laser/LED required for such imaging could exceed the optical system’s thresholds and potentially damage the optical components. Additionally, reflections from bright particles on the microbubble’s surface can degrade the quality of tracking in the vicinity of the bubble, making brightfield the preferred method in these scenarios.

### 2.4. Control system setup

The operational control system, managed by LabVIEW (National Instruments, USA), is essential for the precise execution of the cell removal process. This system manages real-time image acquisition (50 fps was sufficient for this study) and performs image analysis to detect and track compromised cells. Once a damaged cell is identified within the field of view, LabVIEW controls the microbubble’s activation and deactivation based on the cell’s initial lateral position (*y*_cell_) and the desired target position (*y*_target_) downstream in the channel. Although this is the first instance of its application to biological cells, the use of such control systems has been effectively demonstrated for precise particle positioning within microfluidic environments, offering a relevant context for its application in this study (Bakhtiari & Kähler, 2022).

The microbubble generates specific flow patterns that are critical for maneuvering cells. If a cell’s location in the y-direction (*y*_cell_) differs from the desired predefined y-Position (*y*_target_), the system activates the microbubble to initiate streaming flows that move the cell either downward, toward the bubble’s sides (negative *y*-direction), or upward, away from the bubble (positive *y*-direction) to reach the *y*_target_. Downward flows, present both upstream and downstream of the bubble, guide cells toward the negative *y*-direction, while the upward flow directly above the bubble pushes cells away from it. Once the cell reaches the target *y*-position, the microbubble is deactivated to maintain the cell at this level for the remainder of its passage through the channel.

To facilitate this process, a function generator (GW INSTEK AFG-2225) and an amplifier (Krohn-Hite 7500) deliver a predefined electrical signal, tuned to the microbubble’s resonant frequency, to the piezoelectric transducer on the microfluidic chip. The function generator is synchronized with a National Instruments USB-6002 DAQmx data acquisition device, controlled by LabVIEW. This setup allows for precise triggering of the function generator at specific intervals and durations, making the system adaptable to various experimental conditions, including different flow rates, cell sizes, channel dimensions, and microstreaming intensities, as illustrated in Figure 2.

**Figure 2.**
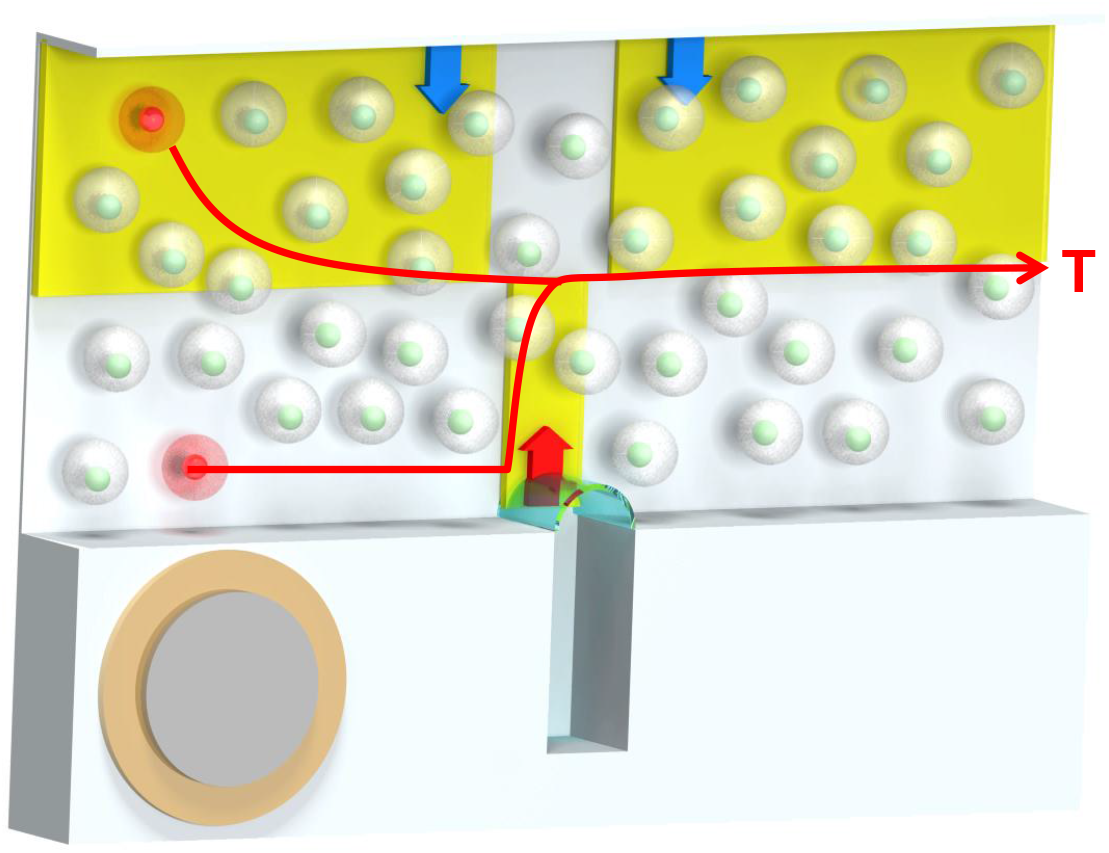
The customized LabVIEW control system manages real-time cell detection, tracking, and precise control of the piezoelectric transducer to guide cells along the red pathways toward the target region (T) for collection in a downstream chamber. The regions of interest (ROIs) for cell detection are highlighted in yellow, with blue arrows indicating the downward flow direction and the red arrow above the bubble showing the upward flow generated when the microbubble is activated.

## 3. Results and Discussion

### 3.1. Automatic Removal of Dead Cells

For the first time, a series of systematic experiments demonstrated the use of an automated cell positioning and removal technique on biological cells. This method autonomously identifies and monitors compromised cells. Once detected, they are removed from the primary fluid using ultrasound-driven microbubble streaming.

In this study, we used a diluted solution of propidium iodide containing white cells (see 2.1.2). The solution had a concentration of 1000 cells per microliter. Around 1 in 1000 cells were damaged. These damaged cells emitted red fluorescence, making them detectable under fluorescent optical microscopy.

The experiments focused on positioning these individual cells within two regions across the width of a 500 μm microchannel. The regions included the upper and bottom parts of the channel. The method allowed cells to be redirected effectively. This system has practical applications, enabling operators to guide individual cells into a designated outlet downstream. Cells could either be collected for analysis or sent to a waste exit, thereby purifying the main fluid.

The experiments used microbubbles with dimensions (*w* = 80 μm and *d* = 0.5 a) and oscillating at a frequency of 18.9 kHz and voltage of 70 v_pp_. Fluid was pumped into the channel at a flow rate of 0.014 μl/s. The bubble’s surface height was controlled by maintaining a constant pressure difference between the inlet and ambient pressure, ensuring no changes in bubble size.

Figure 3 presents the averaged results of image stacks from 14 independent trials each. Figure 3a shows cells targeted to the upper region, while Figure 3b displays those directed to the lower region. To enhance the visibility of compromised cells, shown in black, the images were inverted and thresholded. This minimized the visibility of live cells, which were abundant in the fluid but filtered out in the demonstration results. In both cases, damaged cells were accurately directed to their intended locations—upper in Figure 3a and lower in Figure 3b.

**Figure 3.**
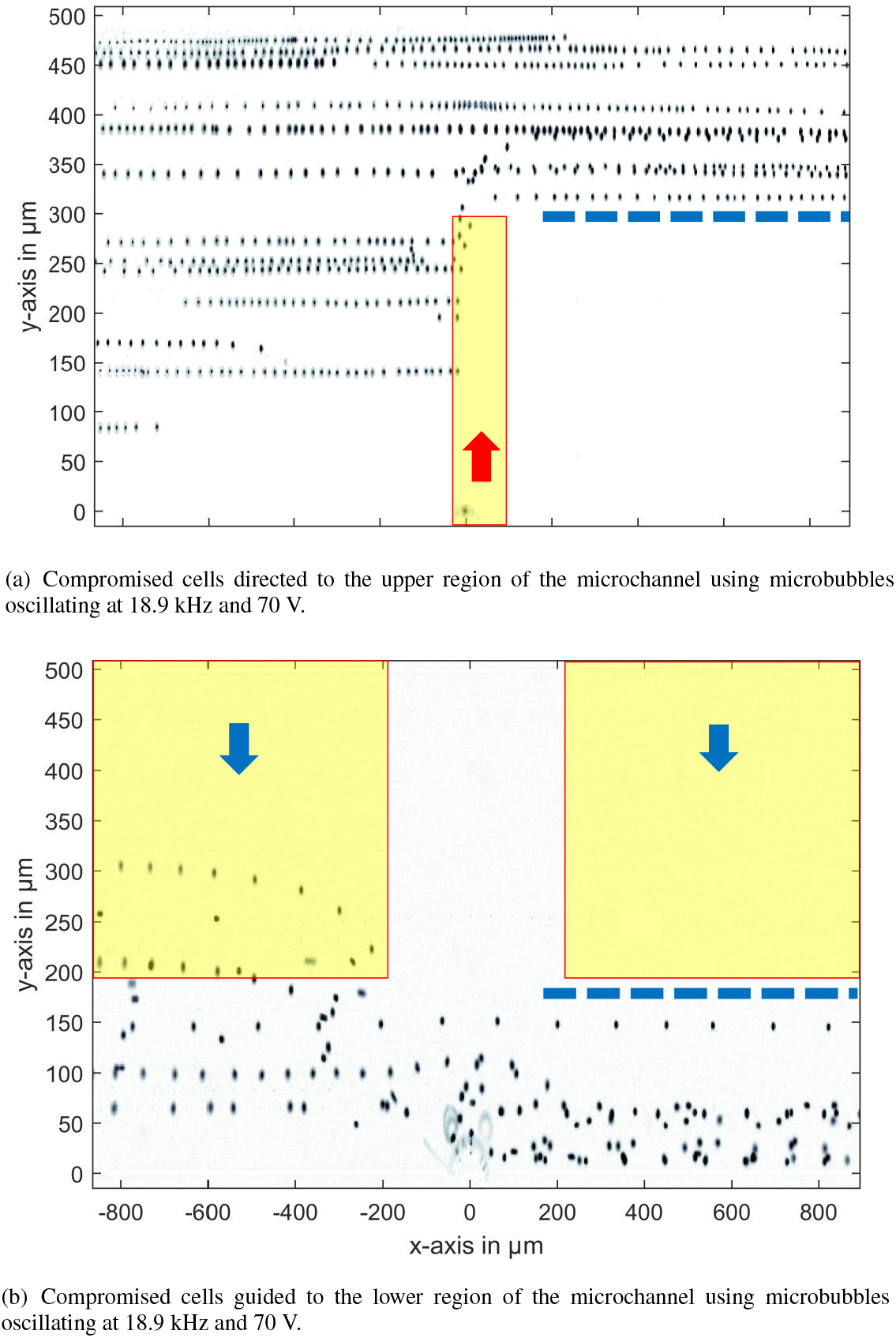
Averaged results from 14 trials each showing the redirection of damaged cells in a 500 μm microchannel using oscillating microbubbles. Images were inverted for enhanced visualization.

Across all trials, the system reliably redirected cells to the desired positions. Notably, although the technique was initially designed for handling single or rare cells, in some cases where multiple cells were present in FOV the system successfully managed numerous cells within the field of view. That demonstrates the robustness and applicability of the approach to more complex scenarios. These findings confirm that the proposed method can autonomously detect, track, and guide biological cells to precise downstream locations, indicating its versatility and potential for broader applications in lab-on-a-chip systems and medical diagnostics.

### 3.2. Shear Stress Induced by an Actuated Microbubble

When investigating cell manipulation via bubble-induced flow, ensuring that cells are not exposed to damaging levels of shear stress is paramount. In this study, we utilized 2 μm buoyant polystyrene particles to measure velocity fields within the vortex flow induced by the microbubble, and based on these measurements, we calculated the shear stress distribution around the bubble (see Figure 4).

**Figure 4.**
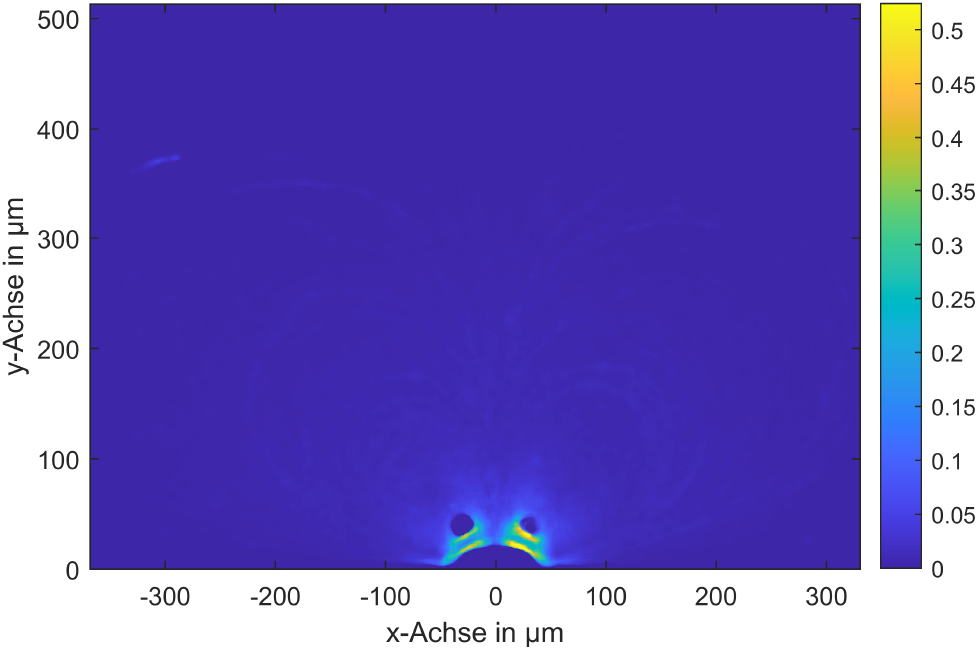
Shear stress due to the actuated bubble flow.

Our measurements revealed that shear stress is highest near the bubble surface, with peak values reaching around 0.5 N / m^2^. This value is significantly lower than the critical shear stress thresholds for most biological cells, such as 150 N / m^2^ for red blood cells (RBCs) or 10 N / m^2^ for white blood cells (WBCs) (Zhao et al., 2022; Espina et al., 2023). Furthermore, the highest shear stress region is highly localized, confined to a very small area near the bubble, and the stress drops rapidly just a few micrometers away from the surface. This rapid decrease in shear stress ensures that biological cells located further from the bubble experience minimal stress, thus significantly reducing the likelihood of cellular damage.

In the core of the induced vortices, velocities—and consequently, shear stresses—could not be measured accurately, as even the small 2 μm tracer particles did not enter these regions. If one assumes that the fluid in the core of the vortex performs a pure rotation, then the shear stress in the vortex core, i.e. the area without measured velocities, can be determined analytically from the actually measured velocities. This calculation shows very low shear stresses, which are 10-fold below the maximum shear stress in the area of the bubble. This, nevertheless, creates some uncertainty in the shear stress characterization at the vortex centers. However, this does not pose a significant problem for cell viability, as biological cells are much larger than the tracer particles and are not expected to be swept into these high-stress areas. The localized nature of the high-shear region, combined with the rapid stress reduction further away from the bubble, ensures that cells remain protected from potential damage while being efficiently manipulated by the microbubble streaming. This data is critical in reinforcing the biocompatibility of ultrasound-driven microbubble streaming, confirming that the system operates well within safe shear stress limits for manipulating biological cells without inducing mechanical damage. For applications where the shear stresses for cells or particles to be selected are particularly critical, these sensitive particles should be moved to the channel side of the bubble. For this particle movement, the particles are only moved through the two larger monitoring areas. Only very low shear stresses act on the particles or cells, as the actuation is only switched on when the particles are located in these areas far away from the bubble.

### 3.3. Monitoring Cell Viability Under Shear Stress Induced by Microbubble Streaming

To build on the previous findings, where the shear stress distribution around the microbubble was mapped (in Figure 4), it was essential to verify these results using live cells to assess the biocompatibility of the induced flows. A critical test was performed in the microchannel under conditions where Poiseuille flow was absent, allowing the effects of microbubble-induced shear stress on cells to be isolated. In this experiment, cells were exposed to the localized shear stress fields generated by the oscillating microbubble for over two hours. To evaluate whether the cells were subjected to damage under these conditions, PI was introduced into the aqueous phase. By using this marker, it was possible to precisely monitor cell viability in real time during the experiment. To minimize the potential for photodegradation of the dye and cellular damage caused by continuous exposure to high-intensity light, the high-power LED was pulsed and synchronized with the camera’s recordings at a rate of 0.1 Hz. The results showed despite the cells being subjected to the shear stress induced by the microbubble for two hours, none of the approximately 40 cells in the field of view displayed any signs of damage. No fluorescence was observed, indicating that none of the cells had compromised membranes—an indicator that they remained undamaged throughout the experiment. Given these findings, it is safe to infer that, under typical microbubble streaming conditions, the likelihood of cell damage is exceedingly low, even during extended exposure periods. This reinforces the viability of microbubble streaming as a reliable and safe method for manipulating cells in microfluidic systems, offering a powerful tool for cell sorting, purification, and positioning without negatively impacting cell health.

## 4. Conclusions and Recommendations

This study effectively demonstrated the viability of using ultrasound-driven microbubbles for precise human cell manipulation in microfluidic environments, while preserving cell integrity. The automated system successfully detected, tracked, and redirected compromised cells, highlighting its potential for applications in lab-on-a-chip systems and medical diagnostics. Additionally, microbubble streaming proved to be a reliable method for removing damaged cells from the primary fluid stream.

Detailed measurements of the shear stress distribution near the microbubble showed that while the highest stress occurred close to the bubble, it decreased rapidly with distance. The peak shear stress was 0.5 N/m^2^, well below the critical limit for most biological cells and confined to a small area near the bubble. This localized stress distribution ensures that cells experience minimal shear stress beyond this region, significantly reducing the risk of cellular damage. Moreover, the study confirmed that cells, being larger than the tracer particles used, are not at risk of being swept into the high-shear zones, ensuring their protection from potential harm.

The monitoring of cell viability under shear stress over extended periods provided further confirmation of the biocompatibility of microbubble streaming. Cells exposed to the bubble-induced flow for over two hours showed no signs of damage, as evidenced by the absence of fluorescent signals from propidium iodide staining. This outcome suggests that the shear stress induced by microbubble streaming is well below the threshold that would compromise cell membrane integrity, even during prolonged exposure.

In summary, this work highlights the biocompatibility and precision of microbubble streaming for non-invasive cell manipulation and sorting. The findings support its broader implementation in biomedical applications, such as automated dead cell removal and selective cell positioning, without negatively impacting cell viability. Future research could explore the use of microbubble streaming in more diverse biological systems and further optimize control mechanisms to enhance precision under varying flow conditions.

## Acknowledgements

We acknowledge financial support by Universität der Bundeswehr München.

## Author Contributions

Amirabas Bakhtiari: conceptualization, methodology, data acquisition and analysis, original draft writing.

Benedikt Schumm: conceptualization, methodology, data acquisition and analysis, original draft writing.

Martin Schönfelder: conceptualization, methodology, review and editing.

Christian J. Kähler: conceptualization, methodology, supervision, review and editing, funding acquisition.

## Data Availability Statement

Raw data are available from the corresponding author Amirabas Bakhtiari.

## Ethical Standards

The research meets all ethical guidelines, including adherence to the legal requirements of the study country.

## References

Armstrong, A. J., Marengo, M. S., Oltean, S., Kemeny, G., Bitting, R. L., Turnbull, J. D., Herold, C. I., Marcom, P. K., George, D. J., & Garcia-Blanco, M. A. (2011). Circulating tumor cells from patients with advanced prostate and breast cancer display both epithelial and mesenchymal markers. Molecular Cancer Research, 9(8), 997–1007.

Baeyens, N., Bandyopadhyay, C., Coon, B. G., Yun, S., Schwartz, M. A., & others. (2016). Endothelial fluid shear stress sensing in vascular health and disease. The Journal of Clinical Investigation, 126(3), 821–828.

Bakhtiari, A., & Kähler, C. J. (2023). Automated microparticle positioning using a pair of ultrasound-actuated microbubbles for microfluidic applications. Microfluidics and Nanofluidics, 27(6), 37.

Bakhtiari, A., & Kähler, C. J. (2024). Enhanced particle separation through ultrasonically-induced microbubble streaming for automated size-selective particle depletion. RSC Advances, 14(4), 2226–2234.

Bakhtiari, A., & Kähler, C. J. (2024). A method to prevent clogging and clustering in microfluidic systems using microbubble streaming. Biomicrofluidics, 18(4).

Bischoff, F. Z., Marquez-Do, D. A., Martinez, D. I., Dang, D., Horne, C., Lewis, D., & Simpson, J. L. (2003). Intact fetal cell isolation from maternal blood: improved isolation using a simple whole blood progenitor cell enrichment approach (RosetteSep™). Clinical Genetics, 63(6), 483–489.

Khine, M., Lau, A., Ionescu-Zanetti, C., Seo, J., & Lee, L. P. (2005). A single cell electroporation chip. Lab on a Chip, 5(1), 38–43.

Koutsiaris, A. G., Tachmitzi, S. V., Batis, N., Kotoula, M. G., Karabatsas, C. H., Tsironi, E., & Chatzoulis, D. Z. (2007). Volume flow and wall shear stress quantification in the human conjunctival capillaries and post-capillary venules in vivo. Biorheology, 44(5-6), 375–386.

Marmottant, P., & Hilgenfeldt, S. (2003). Controlled vesicle deformation and lysis by single oscillating bubbles. Nature, 423(6936), 153–156.

Nan, L., Jiang, Z., & Wei, X. (2014). Emerging microfluidic devices for cell lysis: a review. Lab on a Chip, 14(6), 1060–1073.

Nilsson, J., Evander, M., Hammarström, B., & Laurell, T. (2009). Review of cell and particle trapping in microfluidic systems. Analytica Chimica Acta, 649(2), 141–157.

Riley, N. (2001). Steady streaming. Annual Review of Fluid Mechanics, 33(1), 43–65.

Roux, E., Bougaran, P., Dufourcq, P., & Couffinhal, T. (2020). Fluid shear stress sensing by the endothelial layer. Frontiers in Physiology, 11, 861.

Sheng, W., Ogunwobi, O. O., Chen, T., Zhang, J., George, T. J., Liu, C., & Fan, Z. H. (2014). Capture, release and culture of circulating tumor cells from pancreatic cancer patients using an enhanced mixing chip. Lab on a Chip, 14(1), 89–98.

Shields, C. W., Reyes, C. D., & López, G. P. (2015). Microfluidic cell sorting: a review of the advances in the separation of cells from debulking to rare cell isolation. Lab on a Chip, 15(5), 1230–1249.

Thameem, R., Rallabandi, B., & Hilgenfeldt, S. (2016). Particle migration and sorting in microbubble streaming flows. Biomicrofluidics, 10(1), 014124.

Versluis, M., Goertz, D. E., Palanchon, P., Heitman, I. L., van der Meer, S. M., Dollet, B., de Jong, N., & Lohse, D. (2010). Microbubble shape oscillations excited through ultrasonic parametric driving. Physical Review E, 82(2), 026321.

Volk, A., & Kähler, C. J. (2018). Size control of sessile microbubbles for reproducibly driven acoustic streaming. Physical Review Applied, 9(5), 054015.

Volk, A., & Kähler, C. J. (2018). Density model for aqueous glycerol solutions. Experiments in Fluids, 59(5), 75.

Wang, C., Jalikop, S. V., & Hilgenfeldt, S. (2010). Trapping, focusing and sorting of microparticles through bubble streaming. arXiv preprint, 1010.3290.

Wang, C., Jalikop, S. V., & Hilgenfeldt, S. (2012). Efficient manipulation of microparticles in bubble streaming flows. Biomicrofluidics, 6(1), 012801.

Wang, C., Rallabandi, B., & Hilgenfeldt, S. (2013). Frequency dependence and frequency control of microbubble streaming flows. Physics of Fluids, 25(2), 022002.

